# Single molecule studies characterize the kinetic mechanism of tetrameric p53 binding to different native response elements

**DOI:** 10.1101/2022.06.01.494191

**Authors:** Johannes P. Suwita, Calvin K. Voong, Elina Ly, James A. Goodrich, Jennifer F. Kugel

## Abstract

The transcriptional activator p53 is a tumor suppressor protein that controls cellular pathways important for cell fate decisions, including cell cycle arrest, senescence, and apoptosis. It functions as a tetramer by binding to specific DNA sequences known as response elements (REs) to control transcription via interactions with co-regulatory complexes. Critical for understanding how p53 regulates gene expression is unraveling the fundamental mechanisms by which it binds to REs. Toward this goal we have used an in vitro single molecule fluorescence approach to quantify the dynamic binding of tetrameric p53 to five native REs in real time under equilibrium conditions. We found little evidence of dimer/DNA complexes as intermediates to the formation or dissociation of p53 tetramer/DNA complexes; however, tetramer/DNA complexes can exchange dimers at some REs. Determining rate constants for association and dissociation revealed two kinetically distinguishable populations of tetrameric p53/RE complexes. For the less stable population, the rate constants for dissociation were larger at REs closest to consensus, showing the more favorable binding sequences form the least kinetically stable complexes. Together our real time measurements provide insight into mechanisms with which tetrameric p53 forms complexes on different native REs.

## Introduction

The transcriptional activator and tumor suppressor p53 is one of the most widely studied mammalian proteins. The p53 protein forms a tetramer consisting of four monomers (393 amino acids each) that each contain a core DNA binding domain and oligomerization domain flanked by an acidic N-terminal region and an unstructured C-terminal tail [1,2]. p53 tetramers bind DNA at p53 response elements (REs), which are composed of two 10 bp half site sequences separated by 0–13 bp [3,4]. The half site consensus sequence is RRRCWWGYYY, where R represents A or G, W represents A or T, and Y represents C or T [3,4]. p53 REs are found throughout mammalian genomes, and p53 binding controls the transcription of genes involved in multiple cellular pathways including stress response pathways and cell cycle control [5,6]. Most of the biochemical and structural studies of p53 have been performed with isolated domains, such as the DNA binding domain. These studies have yielded critical insight into understanding how p53 interacts with DNA; and importantly, how the many mutations in the DNA binding domain that are associated with a variety of cancers impair the function of the protein [7,8]. Indeed, mutations that disrupt the ability of p53 to bind DNA are among the most prevalent found in tumors, underscoring the importance of understanding how this protein interacts with its REs [9–11].

To gain insight into the mechanisms by which p53 binds to DNA, we previously studied full-length human p53 tetramers interacting with the RE from the promoter of the human GADD45 gene in real time using in vitro single molecule fluorescence co-localization assays [12]. This enabled us to investigate p53/DNA complexes under equilibrium conditions, which was important for observing binding specificity. In our single molecule studies p53 tetramers readily bound the GADD45 RE but not a randomized RE, whereas in electrophoretic mobility shift assays we found that p53 bound to a randomized DNA with low nM affinity [12]. The single molecule system resolved rapid binding and unbinding events, which revealed that p53/ RE interactions are highly dynamic, with many binding and release events occurring over tens of seconds. Notably, single particle tracking experiments in live cells have also shown that p53 interacts with chromatin with rapid dynamics. In these experiments, two kinetic populations were observed with average residence times of ~3 s and ~0.3 s [13,14].

Rapid interactions with chromatin are not unique to p53. Indeed, live cell single particle tracking studies of many different mammalian transcription factors reveal that, in general, transcription factor binding is very dynamic [15]. As examples, brief chromatin residence times have been measured for Sox2, Oct4, cMyc, CTCF, SRF, and steroid receptors; moreover, many of these studies report two populations with short (<1s) and somewhat longer (several seconds) residence times [16–20]. The interactions of transcription factors with chromatin that occur on the order of seconds are thought to represent functional interactions with the genome; indeed, studies have shown they correlate with transcriptional activity [13]. While single particle tracking approaches enable measurements of transcription factors dynamically binding chromatin in live cells, they are not able to distinguish binding to a specific RE. Hence, in vitro kinetic measurements are required to distinguish how binding rates differ between native REs.

To evaluate the relationship between p53 binding kinetics and the sequence of REs, we determined how natural variation in the p53 RE sequence affects the kinetics with which p53 tetramers associate with and dissociate from DNA. We used an in vitro single molecule fluorescence system to quantify the kinetic parameters and mechanism of p53 binding to five native REs (PTEN, GADD45, MDM2, p21, and PUMA) that differ from the consensus sequence by variable amounts. On all REs, we found little evidence of dimer/DNA complexes en route to binding or dissociation of p53 tetramers. On the p21 and GADD45 REs in particular, DNA-bound tetramers exhibited exchange of dimers. Rate constants for binding and dissociation of p53 tetramers were measured on all REs. The data revealed two kinetically distinguishable populations of p53/DNA complexes, consistent with prior models [12]. Unexpectedly, the measured rate constants for dissociation of the less stable complexes were larger (i.e. more rapid release) at REs closest to consensus, showing the most favorable binding sequence forms the least stable complexes.

## Materials and Methods

### DNA constructs and AF647-p53

Oligonucleotides were ordered HPLC purified from Integrated DNA Technologies. All forward oligonucleotides had a 5’ biotin tag attached to a 24 nt single stranded linker: 5’-CGCGTTCATGGTAGAGTCGTGGAC-3’. All reverse oligonucleotides had a 5’ AF647 dye. The sequences of the oligonucleotides were (5’ to 3’):

PTEN forward, TAGAGCGAGCAAGCCCGGGCATGCTCGCGTCG; PTEN reverse,

CGACGCGAGCATGCCCGGGCTTGCTCGCTCTA;

GADD45 forward, TAGAGCGAACATGTCTAAGCATGCTGGCGTCG; GADD45 reverse,

CGACGCCAGCATGCTTAGACATGTTCGCTCTA; p21 forward,

TAGAGCGAACATGTCCCAACATGTTGGCGTCG; p21 reverse,

CGACGCCAACATGTTGGGACATGTTCGCTCTA-3’; MDM2 forward,

TAGAGCGGTCAAGTTCAGACACGTTCGCGTCG; MDM2 reverse,

CGACGCGAACGTGTCTGAACTTGACCGCTCTA; PUMA forward,

TAGAGCCTGCAAGTCCTGACTTGTCCGCGTCG; PUMA reverse,

CGACGCGGACAAGTCAGGACTTGCAGGCTCTA; Randomized forward,

CGTTCCTAATCGGGTAGGGGCTGACACGAATA; Randomized reverse,

TATTCGTGTCAGCCCCTACCCGATTAGGAACG.

Forward and reverse oligonucleotides were annealed in 1X annealing buffer (20 mM Tris pH 7.9, 2 mM MgCl2, 50 mM KCl), with 4 μM reverse oligo and 1 μM forward oligo by incubating at 95 °C for 10 min followed by slow cooling to room temperature. Annealed oligos were gel purified after resolving on a 7% native polyacrylamide gel containing 0.5X TBE. The gel was stained (0.01% SYBR^®^ Gold, 0.5X TBE) and the desired band was cut out and transferred to a 1.5 mL reaction tube. 380 μL of TE Low (10 mM Tris pH 7.9, 0.1 mM EDTA) and 20 μL 1 M KCl were added to the tube, the gel piece was crushed, and the tube was nutated over night at 4 °C. The gel pieces were pelleted via centrifugation and the supernatant was ethanol precipitated.

AF647-p53 was made as previously described [12]. Briefly, full length human p53 containing an N-terminal His tag and a C-terminal SNAP tag was expressed in SF9 insect cells using a recombinant baculovirus. The protein was purified from cellular extracts using HisPur Ni–NTA resin (Thermo Scientific). The purified protein was labeled with SNAP-Surface Alexa Fluor647 dye (New England Biolabs). The fractional dye labeling and DNA binding activity were assessed as previously described [12].

### Single molecule binding assays

Microscope slides and coverslips were prepared as described [21]. In brief, slides and coverslips were cleaned in a water bath sonicator in multiple steps with 1% alconox, 100% ethanol, 1 M KOH and 100% methanol. Then they were functionalized with aminosilane, PEGylated with biotin-PEG/mPEG, and the flow chambers were assembled. All buffers and solutions needed for the assays were prepared as described [12,21].

To perform the single molecule binding assays, the flow chamber was washed twice with 200 μL ddH_2_O then twice with 200 μL 1X buffer (25 mM Tris pH 7.9, 50 mM KCl, 5 mM MgCl_2_, 10% glycerol, 0.05 mg/mL BSA, 1 mM DTT, 0.1% NP-40). A streptavidin solution (50 μL of 0.2 mg/mL streptavidin, 0.8 mg/mL BSA diluted in 1X buffer) was flowed into the chamber. After an incubation of 5 min, the unbound streptavidin was removed by washing twice with 200 μL 1X buffer. Next, 100 μL of 10 pM DNA in 1X buffer was flowed into the chamber and incubated for 10 min. Unbound DNA was removed by washing twice with 200 μL 1X buffer. Then, 100 μL of imaging buffer (1.02 mg/mL glucose oxidase, 0.04 mg/mL catalase, 0.83% D-glucose, 3.04 mM Trolox diluted in 1X buffer) was flowed in and DNA-only emission movies were collected over four regions of the slide using a piezo nanopositioning stage. The flow chamber was washed again with 200 μL of 1X buffer. AF647-p53 (1.0 nM, monomeric concentration) diluted in imaging buffer containing 1.25 mg/mL BSA was flowed into the chamber and DNA+p53 emission movies were collected over the same four regions.

Emission movies were collected with an objective based TIRF microscope (1.49 NA immersion objective, Nikon TE-2000U microscope) and an Andor iXon Life 897 EMCCD camera using the NIS Elements Software. AF647 fluorophores were excited with a 635 nm laser. The laser intensity was set between 115 mA and 125 mA. DNA+p53 movies were collected for 1000-2000 frames (frame rates of 60 ms, 200 ms or 600 ms); the corresponding DNA-only movies were collected for 100 frames. A neutral density filter was used to reduce photobleaching during long measurements with an exposure of 200 ms or 600 ms.

### Analysis of single molecule data

Emission movies of DNA+p53 were analyzed as described using in-house software written in IDL [12]. Spots in the DNA-only movie and the DNA+p53 movie from the same region of a slide were computationally colocalized. First, fluorescent spots with an intensity within the user-defined range were identified in the summed image of each movie individually. Then, if the distance between a spot in the DNA-only movie and a spot in the DNA+p53 movie was ≤ 1-3 pixels, the two spots were considered a potential spot pair. Each potential spot pair was manually visualized and evaluated. A spot pair was rejected if, for example, two spots in close proximity were identified as one spot.

States (unbound and bound DNA) and state changes were computationally determined from the emission intensity over time of a spot and manually visualized and reviewed. For each spot, the emission intensity in the DNA-only movie was defined as the intensity of one AF647 dye (1N). This value was used to classify the oligomeric state of p53 for bound states at that spot in the DNA+p53 movie. For example, the state of the spot in the DNA+p53 movie was classified as unbound if the emission intensity was 1N, dimer-bound if it was 3N, and tetramer-bound if it was 4N or 5N. The number of states, binding/unbinding transitions, and dwell times were compiled across each region and for multiple slides (biological replicates) for each RE. To determine rate constants, bound dwell times (unbound to tetramer-bound to unbound) and unbound dwell times (tetramer-bound to unbound to tetramer-bound and dimer-bound to unbound to tetramer-bound) from individual replicates were combined. To ensure the number of states from movies with different frame rates was approximately equal, the area of the slide region quantified was adjusted. The dwell times were plotted as cumulative sums, and non-linear regression of single and double exponential equations was performed using Solver in Microsoft Excel. Half times for binding and unbinding were calculated using t_1/2_ = ln2/k. 95% confidence intervals were obtained using GraphPad Prism 9.

## Results

### A single molecule system for studying p53 tetramers dynamically binding to DNA

We previously developed a single molecule fluorescence system for studying the binding of tetrameric p53 to DNA using a TIRF (total internal reflection fluorescence) microscope [12]. A key component of this system is fluorescently labeled full length recombinant human p53, which is expressed in insect cells with an N-terminal His tag and a C-terminal SNAP tag for labeling (p53-SNAP). After purification, p53-SNAP is labeled with AlexaFluor647 (AF647-p53). AF647-p53 is fully active for DNA binding when compared to wild-type p53 [12]. The dye conjugation is near 100% as assessed by LC MS/MS, and ~70% of the dyes appear photoactive [12].

An overview of the single molecule system used to study DNA binding by AF647-p53 is shown in Fig 1A. First, AF647-labeled DNA containing a p53 RE is immobilized on the surface of a slide chamber via a biotin tag. We then image the immobilized AF647 DNA molecules in 4 regions to register their locations and emission intensity using a TIRF microscope (i.e. DNA-only movie). AF647-p53 is then flowed into the chamber and we image the same four regions over time to monitor p53 binding events (i.e. DNA+p53 movie). For each region, spots of AF647 emission from the DNA+p53 movie are co-localized with spots of AF647 emission from the DNA-only movie. To identify and quantify p53 tetramer binding and dissociation events, the fluorescence emission is quantified over time for each spot pair. The fluorescence intensity of the AF647 emission from a given spot in the DNA-only movie is used to obtain a baseline intensity for a single red dye at that position on the surface. Using this baseline, the number of dye molecules associated with each AF647-p53 binding and dissociation event is calculated, which yields the oligomeric state of p53 for each binding and dissociation event. Therefore the dynamic interaction of p53 tetramers with a RE can be studied in real time under equilibrium conditions.

**Fig 1.**
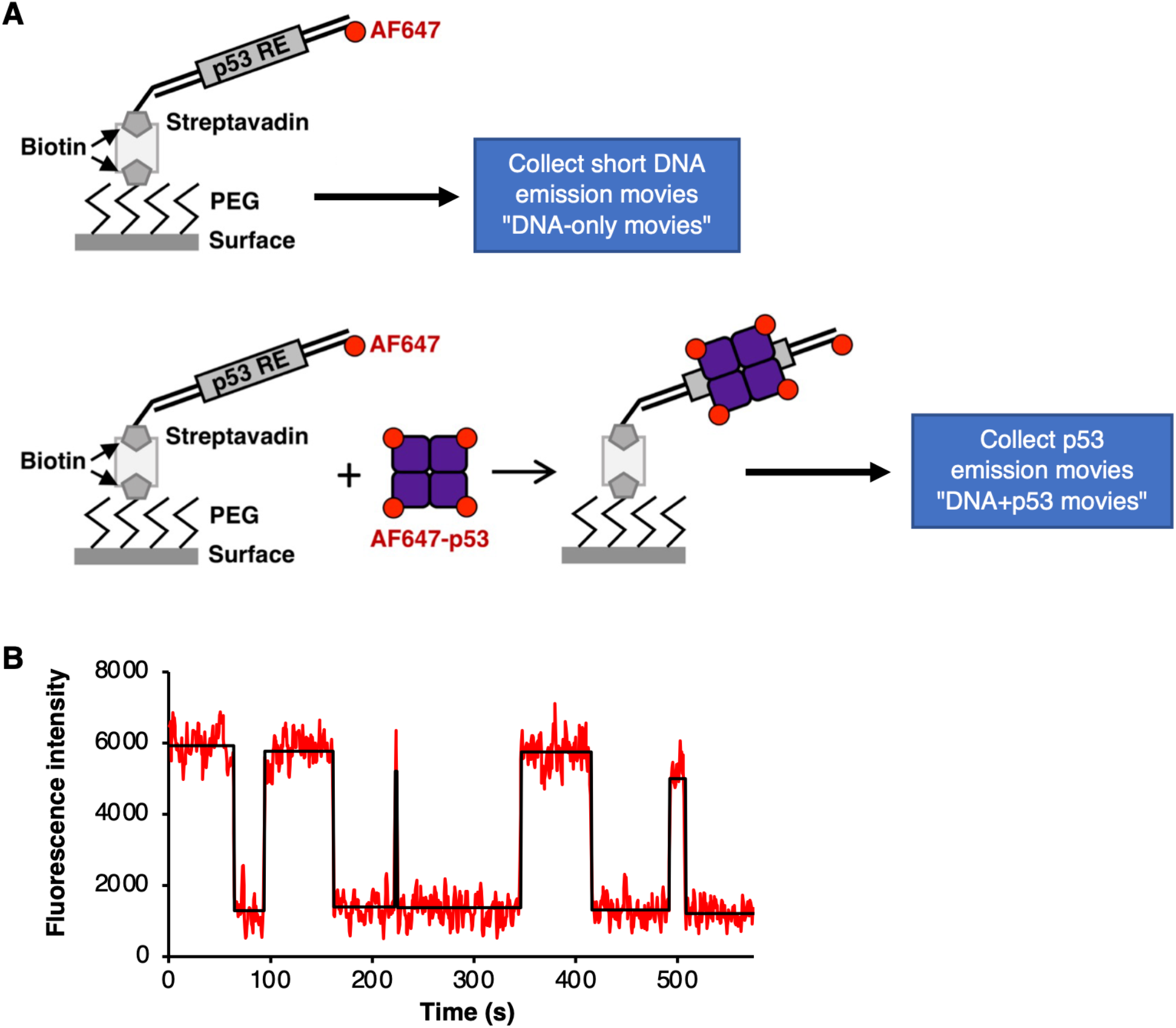
A single molecule fluorescence assay to study dynamic DNA binding by tetrameric p53. **(A)** Illustration of the single molecule assay used. Red emission movies were collected of biotinylated AF647 labeled DNA immobilized on the streptavidin derivatized surface. A second red emission movie (DNA + p53 movie) was collected over the same region after AF647-p53 was flowed into the slide chamber. p53 biding events on immobilized DNA were identified by co-localization of molecules in the DNA only data and the DNA + p53 data. Emission intensity traces over time were generated for each spot pair, and were used to calculate the base intensity value for a single dye on the DNA and the number of dyes (i.e. oligomeric state) of p53 bound to DNA. Fig is adapted from [12]. **(B)** Representative AF647 emission data from a single DNA molecule with five tetrameric p53 binding events. For this movie, emission data were collected every 600 ms for 958 frames. The immobilized DNA contained the PUMA RE, and the p53 concentration was 1.0 nM (monomer). For purposes of display, the signals were smoothed by averaging three adjacent time frames.

Fig 1B shows representative data for the dynamic binding of p53 at a single DNA molecule containing the PUMA RE over a 575 s movie. The plot of fluorescence intensity over time shows that at the beginning of the movie tetrameric p53 is bound to the DNA, then at ~65 s this molecule of p53 dissociated from the DNA. Throughout the remainder of the movie, four other p53 tetramers bound and dissociated from this same DNA molecule. Each change in fluorescence intensity (i.e. each increase or decrease) in the plot was approximately 4 times the intensity of the DNA molecule, consistent with each event involving binding or dissociation of tetrameric p53. We focused our kinetic measurements on tetrameric binding and dissociation events that showed a change of three or four dye intensity units, which accounts for the fact that only ~70% of the dye molecules are fluorescently active (i.e. some tetramers have only three active dyes). We did not consider events in which two active dyes bound then released from free DNA since it could not be distinguished whether this was due to a bona fide dimer binding or the binding of a tetramer with two dark dyes. Limiting our analysis to tetrameric p53 also ensured that dissociation events were not due to photobleaching or photoblinking, which would not occur simultaneously for the three or four active AF647 dyes on a p53 tetramer.

### p53 tetramers show concerted binding and dissociation on native REs, with some exchange of dimers in tetramer/DNA complexes

We were interested in determining how natural variability in the sequence of the p53 RE affects the mechanism and dynamics of binding by p53 tetramers. To study this we chose five native REs that are bound by p53 in cells and regulate transcription of the corresponding mRNA gene. Each of the native REs has no additional basepairs between the two half sites and all half sites have the most highly conserved C and G nucleotides at positions 4 and 7, respectively (Fig 2A). The PTEN RE (named according to the gene it regulates) matches the p53 consensus. The GADD45 RE, which we previously studied, differs from consensus at only one position. The MDM2 and p21 REs differ from consensus at two positions, and the PUMA RE differs from consensus at three positions. We also designed a partially randomized DNA with only half of the basepairs matching the p53 RE consensus sequence.

**Fig 2.**
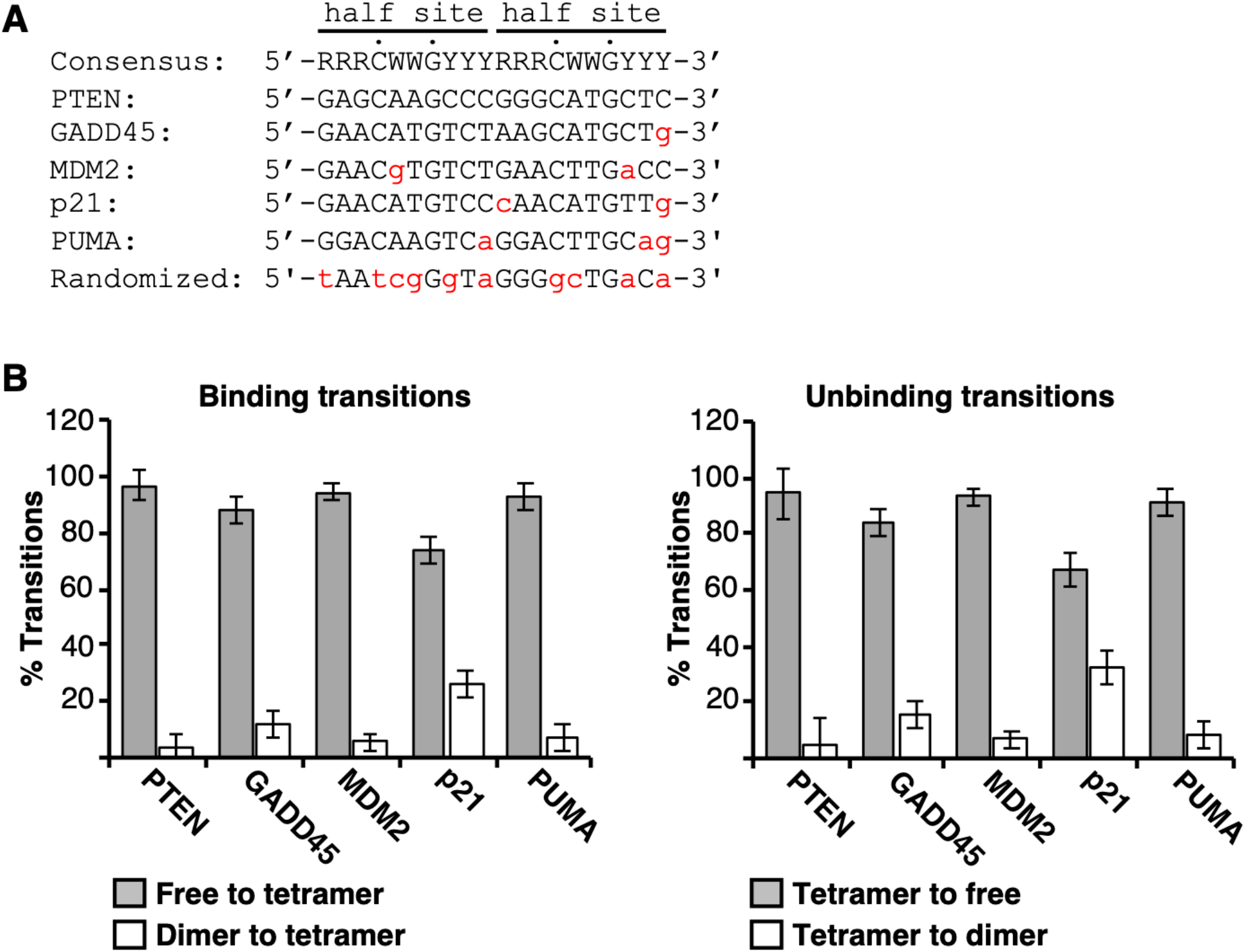
p53 tetramers dynamically associate with five native REs. **(A)** The five native RE sequences and randomized sequence used in the single molecule experiments in comparison to the p53 RE consensus sequence. Positions that are not consensus are shown in red lowercase letters. The two half sites are indicated, and the highly conserved C and G residues in each half site are noted with dots. R represents A or G, W represents A or T, and Y represents C or T. **(B)** p53 tetramer/DNA complexes can undergo dimer exchange. Plotted is the percentage of transitions that were tetramers (e.g. gain/loss of 3 or 4 dyes) or dimers (e.g. gain/loss of 2 dyes when transitioning to or from a tetrameric p53/DNA complex). The errors are the standard deviations of the measurements for the GADD45, MDM2, p21, and PUMA elements (n = 6, 3, 4, and 3 respectively); the error for PTEN is the range of two measurements. The ten pairwise comparisons of free/tetramer transitions (gray bars) to dimer/tetramer transitions (white bars) are statistically different, with p values ≤ 0.005 using an unpaired two-sided t test.

We performed multiple single molecule experiments with each native RE separately immobilized and quantified hundreds of tetrameric p53 DNA binding and dissociation events (i.e. those showing three or four dye units of intensity change). We also performed controls with no DNA on the surface and with the immobilized partially randomized DNA (sequence shown in Fig 2A). In these two conditions we observed very few (less than 10) binding and unbinding events compared to hundreds on the REs. Hence p53 tetramers did not dynamically bind to the slide surface in general or to a DNA with a sequence that is far away from consensus.

On the five REs, for each binding event we evaluated whether tetramer/DNA complexes assembled or disassembled through an intermediate dimer/DNA complex. Because each event began or ended with a tetramer, this analysis is legitimate despite ~70% of dyes being active. We counted each binding and dissociation event that involved the gain or loss of two dye molecules (dimer) en route to the final bound (tetrameric) or unbound (free DNA) state. We then compared this to the number of binding events involving the addition or release of 3 or 4 dye molecules (tetramer). We found that on all REs less than 2% of tetramer/DNA complexes formed via a dimer/DNA intermediate, and less than 5% dissociated via a dimer/DNA intermediate. Therefore, our data support concerted binding and release of p53 tetramers on native REs.

We then investigated whether tetramer/DNA complexes exhibited the exchange of p53 dimers (i.e. dynamic release and rebinding of p53 dimers). To do so, we calculated the frequency of dimer-bound states going to tetramer-bound states compared to the frequency of free DNA going to a tetramer-bound state (Fig 2B, left). We also calculated the frequency of dimers dissociating from tetramer/DNA complexes versus tetramers dissociating to result in free DNA (Fig 2B, right). On three of the five REs (PTEN, MDM2, PUMA), transitions between dimers and tetramers occurred infrequently (the white bars). Only 3-7% of the binding events and 5-8% of the unbinding events involved changes equivalent to two dye molecules. By contrast, on the p21 and GADD45 REs we observed a greater frequency of changes equivalent to 2 dye molecules, with 27% and 12% of binding events, and 33% and 16% of dissociation events, respectively, involving a dimer. Further analyses of the p21 and GADD45 data revealed that ~85% of the binding events and ~85-95% of the release events involved a dimer exchanging on a tetramer/DNA complex. Together our data show that p53 tetramers bound to the p21 and GADD45 REs exhibit release and re-binding of dimers with a greater frequency than on other REs.

### p53 tetramers bind to REs in one or two kinetically distinct steps

We next asked whether there were differences in the kinetics of p53 tetramer binding and release with the five different REs. We began by studying the association of p53 tetramers with the REs. To do so we measured hundreds of unbound dwell times (i.e. the time between two binding events) on each DNA to determine observed rate constants for association. For each RE, the unbound dwell times were plotted as cumulative sums and fit with exponential equations to yield observed rate constants (Fig 3). Data collected on the GADD45, MDM2, and PUMA REs (panels B, C, and E) fit well with a single exponential equation, giving one rate constant for binding. By contrast, data from the PTEN and p21 REs were best fit by a double exponential equation, yielding two forward rate constants (plots of the residuals for fitting the PTEN and p21 data sets with single and double exponential equations are shown in S1 Fig). There is no clear relationship between the RE similarity to consensus and whether the rates of association were best fit by one or two rate constants.

**Fig 3.**
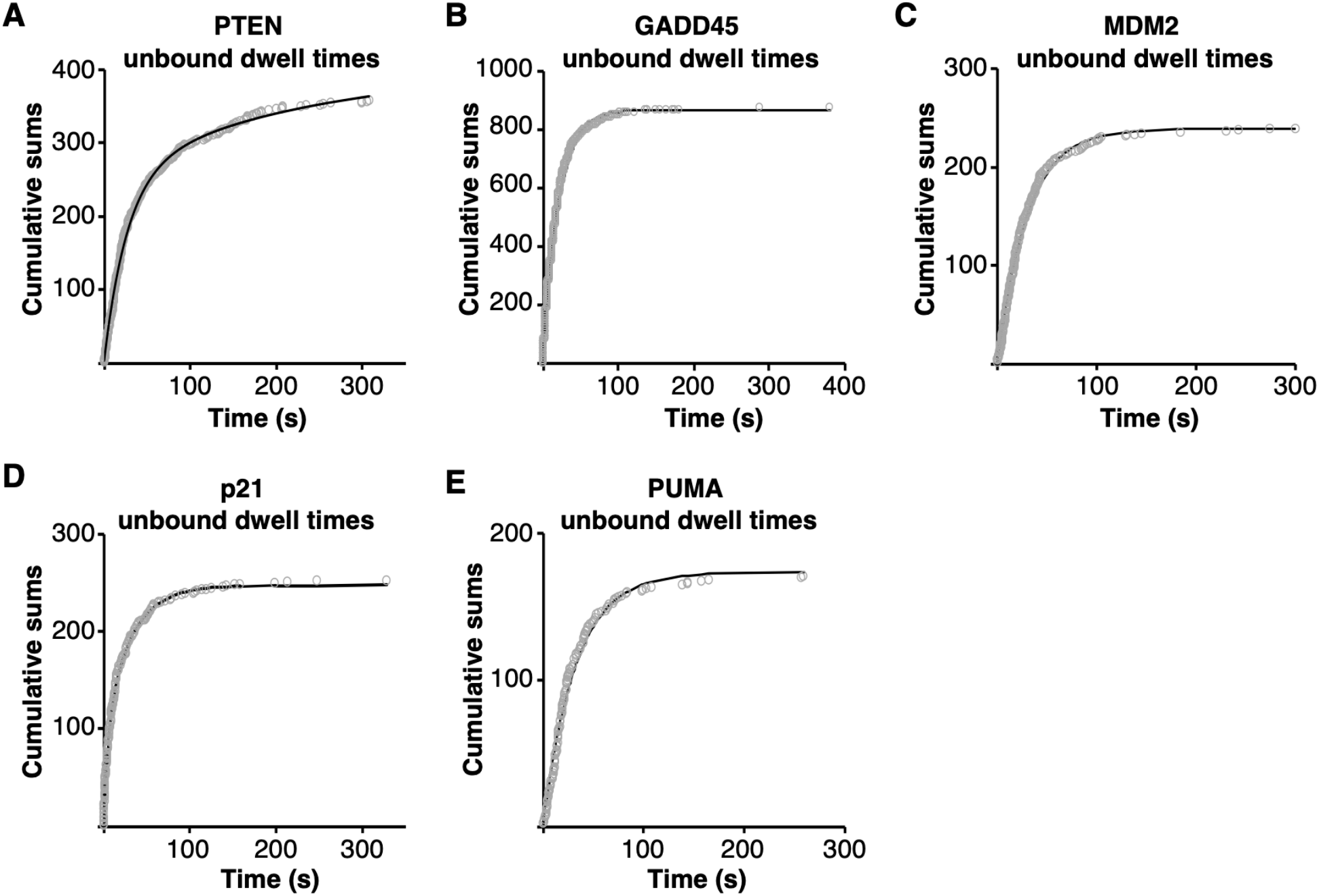
Under equilibrium conditions tetrameric p53 dynamically associated with each of the five REs. Unbound dwell times were plotted as cumulative sums over time and fit with a single or double exponential equation as indicated in the panel legends. The values of the rate constants and 95% confidence intervals from the curve fits are in Table 1. **(A)** PTEN RE, 357 unbound dwell times fit with a double exponential equation. **(B)** GADD45 RE, 870 unbound dwell times fit with a single exponential equation. **(C)** MDM2 RE, 239 unbound dwell times fit with a single exponential equation. **(D)** p21 RE, 252 unbound dwell times fit with a double exponential. **(E)** PUMA RE, 170 unbound dwell times fit with a single exponential.

The values of the association rate constants are shown in Table 1, along with 95% confidence intervals. They are presented as observed first order rate constants (k_on(obs)_, s^-1^) because the measurements were made at a single concentration of p53. The observed rate constants are strikingly similar to one another with the following two exceptions. The larger of the rate constants for the p21 RE is ~6-fold greater than the other k_on(obs)_ values, and the smaller of the rate constants for the PTEN RE is ~8-fold smaller. Thus the REs with two kinetically distinct populations show one population with very rapid binding (p21) and one population with very slow binding (PTEN). In general, there is no correlation between RE sequence or similarity to consensus and the rate constants for association.

**Table 1.** The kinetic constants for tetrameric p53 binding to five native REs.

| DNA | On rate constants |  |  | Off rate constants |  |  |
| --- | --- | --- | --- | --- | --- | --- |
| | $k_{\text{on(obs)}} (\text{s}^{-1})$ | 95% CI | $t_{1/2} (\text{s})$ | $k_{\text{off}} (\text{s}^{-1})$ | 95% CI | $t_{1/2} (\text{s})$ |
| PTEN | 0.037 | 0.036–0.039 | 18.7 | 0.685 | 0.670–0.699 | 1.01 |
|  | 0.004 | 0.002–0.006 | 173 | 0.065 | 0.062–0.066 | 10.6 |
| GADD45 | 0.0479 | 0.0475–0.0483 | 14.47 | 0.49 | 0.48–0.50 | 1.4 |
|  |  |  |  | 0.079 | 0.077–0.081 | 8.77 |
| MDM2 | 0.033 | 0.032–0.034 | 21 | 0.42 | 0.39–0.46 | 1.7 |
|  |  |  |  | 0.12 | 0.10–0.14 | 5.8 |
| p21 | 0.19 | 0.17–0.21 | 3.7 | 0.286 | 0.277–0.298 | 2.42 |
|  | 0.033 | 0.031–0.036 | 21 | 0.02 | 0.01–0.03 | 30 |
| PUMA | 0.031 | 0.030–0.032 | 22.4 | 0.19 | 0.16–0.22 | 3.7 |
|  |  |  |  | 0.05 | 0.04–0.06 | 14 |

### p53 tetramers dissociate via two kinetically distinct steps, with one step being faster from REs that are closer to the consensus sequence

To determine observed rate constants for dissociation of p53 tetramers from each of the five REs, we measured hundreds of bound dwell times for tetrameric p53/DNA complexes. We included only those that formed from free DNA and dissociated to free DNA (i.e. we did not quantify tetramer/DNA complexes that were exchanging dimers, see Fig 2). For each RE the bound dwell times, plotted as cumulative sums, were fit best using a double exponential equation (Fig 4A-E). The plots of the residuals from the single and double exponential fits are shown in S2 Fig. Hence the bound dwell times yield two rate constants for dissociation (k_off1_ and k_off2_, s^-1^) for each RE, which are shown in Table 1, along with 95% confidence intervals. We conclude two kinetically distinct complexes exist on each RE, consistent with our earlier findings on the GADD45 RE [12]. For each RE, the faster dissociating population has a half-time on the order of 1-4 s, while the slower population dissociates ~4-12-fold more slowly (Table 1). The most striking observation regarding the rate constants for dissociation is that the koff1 values (reflecting the least stable population of p53/DNA complexes) increase as the RE sequence moves toward the consensus sequence (Fig 4F). This indicates that for the less stable population of p53/DNA complexes, tetramers dissociate faster from the more favorable binding sequences.

**Fig 4.**
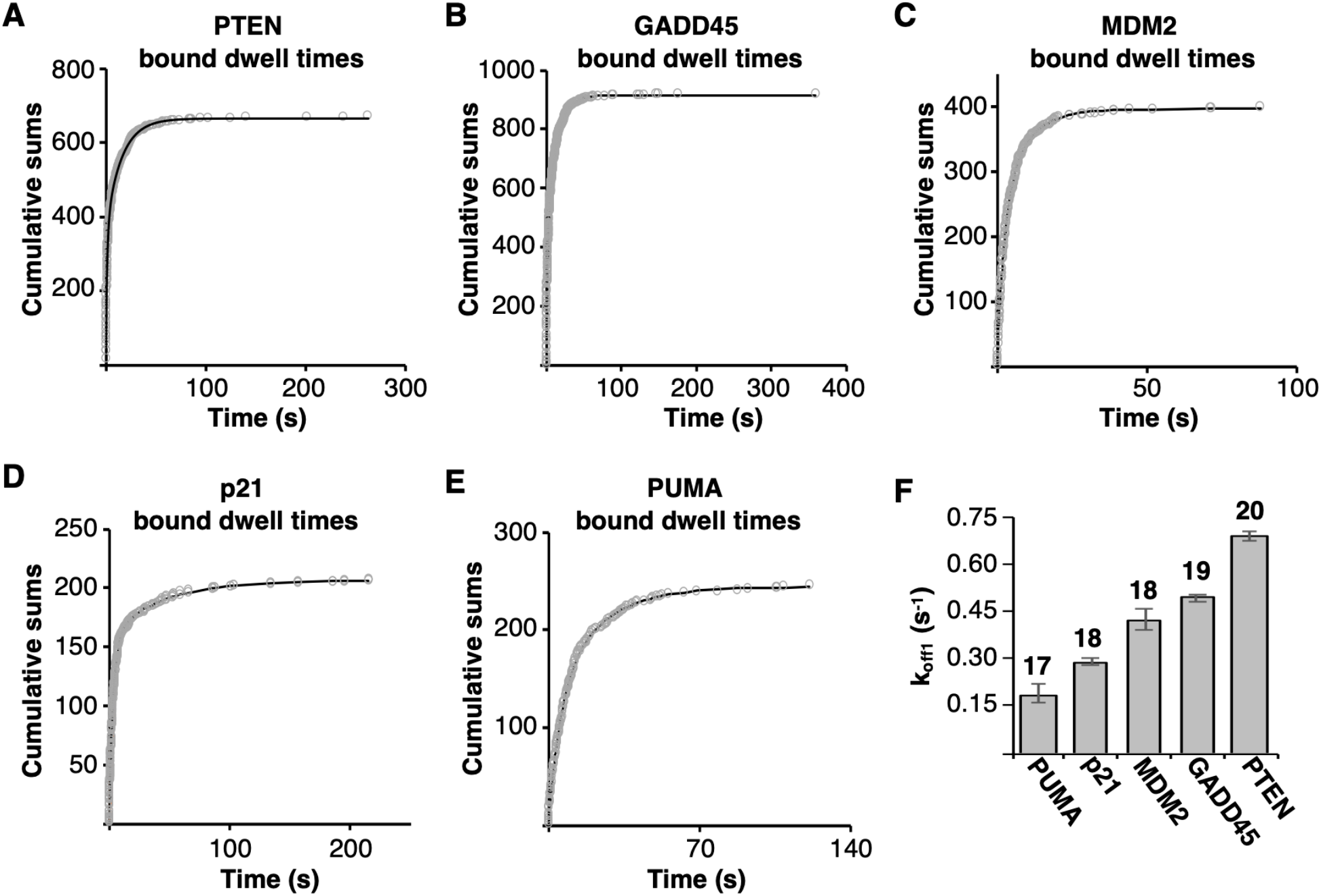
Under equilibrium conditions tetrameric p53 dissociated from each of the five REs in two kinetically distinct steps. Bound dwell times were plotted as cumulative sums over time and fit with a double exponential equation. The rate constants and 95% confidence intervals are shown in Table 1. **(A)** PTEN RE, 671 bound dwell times plotted. **(B)** GADD45 RE, 922 bound dwell times plotted. **(C)** MDM2 RE, 399 bound dwell times plotted. **(D)** p21 RE, 207 bound dwell times plotted. **(E)** PUMA RE, 246 bound dwell times plotted. **(F)** For the less stable p53/ DNA complex, p53 tetramers dissociate more rapidly as the RE sequence becomes more similar to the consensus sequence. Plotted are koff1 values for the five REs, which increase in similarity to consensus from left to right. Above each bar is the number of bp in the RE that match the 20 bp consensus sequence. The error bars are 95% confidence intervals.

## Discussion

We studied the dynamic binding of p53 tetramers to five native REs with variable amounts of divergence from the p53 RE consensus sequence using single molecule fluorescence microscopy. At all REs we found that association and dissociation of tetramers with free DNA is largely concerted, with minimal dimer/DNA intermediates observed, although dimers could exchange at tetramer/DNA complexes on some REs. Real-time kinetic measurements enabled us to determine rate constants governing association and dissociation. Tetrameric p53 bound to DNA in one or two kinetically distinguishable populations, depending on the RE, with no clear relationship between sequence and rate. Tetramers dissociated from all REs in two kinetic populations. Moreover, the rate constants describing the less stable population increased as the sequence of the RE became closer to consensus. In other words, for these complexes similarity to consensus inversely correlates with kinetic stability. Together our studies provide insight into the mechanisms by which p53/RE complexes assemble and disassemble.

Our experimental system uniquely allows us to follow the pathways by which tetrameric p53/RE complexes assemble and disassemble. We found that dimer/DNA complexes infrequently form as intermediates en route to assembly or disassembly of tetramer/DNA complexes (< 5% of the time). This is consistent with prior ensemble biochemical approaches showing that binding of p53 tetramers to REs is concerted [22]. Interestingly, our data also showed that tetramer/RE complexes can exchange dimers (i.e. they show release and re-binding of dimers) at the p21 and GADD45 REs. To our knowledge this exchange has not been reported, which is perhaps not surprising since ensemble systems would be unlikely to allow the exchange to be observed. We do not yet understand why dimer exchange occurs more frequently at p21 and GADD45 compared to the other REs. Our investigations of transitions involving dimer/DNA complexes suggest the possibility that there is an inherent difference in dimer/RE complexes depending on their origin (i.e. whether dimer/DNA complexes arise from binding free DNA or arise from a tetramer/DNA complex losing a dimer). Differences between dimer/RE complexes depending on their origin could be conformational, which will require future investigations to unravel.

Our data show that the tetrameric p53/DNA complexes that form at all five REs exist in minimally two kinetic populations. Specifically, the bound dwell time data yielded two dissociation rate constants, which is consistent with our previous finding for the GADD45 RE [12]. We measured a single rate constant for association at three of the five REs. Our data, in conjunction with existing literature, support a two-step model for binding:

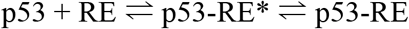

In this model, a p53 tetramer initially binds to the RE to form a relatively unstable complex (p53-RE*) that then undergoes a conformational change to form a more stable complex (p53-RE). Other studies have proposed a two-step binding mechanism that involves an initial binding interaction followed by a conformational change [23–26]. It has been proposed that this induced-fit mechanism of binding allows the selectivity of RE binding to be more dependent on the off-rate kinetics than on the on-rate kinetics [26]. This is interesting given that we found a relationship between the sequence of the REs and the k_off_ values, and not the k_on_ values, for what we believe is the initial p53/DNA interaction. Arriving at a molecular model for the nature of the conformational change that stabilizes the tetrameric p53-RE complex will require additional investigations. In prior work we proposed that the initial recognition and binding event to form the less stable p53-RE* complex primarily involves contacts with one half site, which is followed by interactions with the second half site that stabilize the complex [12].

Of the rate constants measured here, only one showed a clear relationship to sequence. Namely, the larger of the two dissociation rate constants at each RE increased as the RE became more similar to consensus (Fig 4F). In our model, koff1 reflects the stability of the initial p53/RE interaction; hence, we conclude that similarity to consensus correlates with decreased stability of p53-RE* complexes. This was unexpected since increased rates of dissociation can lead to decreased affinity; in particular because we measured similar rates of association across most the REs. Interestingly, the unstructured C-terminal domain of p53 has been shown to facilitate p53 binding to REs that are more divergent from the consensus sequence, leading to the proposal that the C-terminal domain impacts the kinetic stability of complexes [27]. Additional studies will be required to fully explore the relationship between dissociation rates and sequence selectivity for p53, the role of the C-terminal domain, and to determine if transcription factors other than p53 have evolved to favor dynamic interaction with their consensus sequences.

At the majority of REs we measured a single rate constant for association, consistent with the second step in our kinetic model involving an isomerization event. Our data are also consistent with a model in which p53 tetramers form in solution and then bind DNA, since we found little evidence that tetramer/DNA complexes assemble on free DNA via a dimer/DNA intermediate. Given the concentration of p53 flowed into the slide chambers (1 nM monomeric) and the published K_D_ for tetramerization [28], it is possible that only a fraction of the p53 was tetrameric. Therefore, the initial rate of association with DNA could be set by the rate of formation of tetramers. On the p21 and PTEN REs we measured two k_on_ values, suggesting there is a branch, or second entry point into the binding pathway at these REs. For example, p53 could also bind the RE in the more stable conformation (p53-RE) directly. It is possible that a branched binding path is not unique to p21 and PTEN, but does not occur with an experimentally distinguishable on-rate at the other REs.

The kinetic measurements we report here for p53/DNA complexes in vitro are consistent with measurements made for p53/chromatin interactions in live cells [13,14], with interactions lasting only seconds in both cases. This rapid kinetic profile for a transcription factor binding to chromatin in cells is not unique to p53. Short residence times on chromatin have been measured for Sox2, Oct4, cMyc, CTCF, SRF, and steroid receptors; moreover, many of these studies report two kinetic populations with very rapid (<1s) and slightly longer (several seconds) residence times [16–20]. Hence single molecule approaches that enable real-time kinetic measurements (in cells and in vitro) have led to an evolution of the traditional regulatory models of stable binding between transcription factors and their REs. Future in vitro single molecule measurements and live cell imaging can be used to further refine dynamic models for transcriptional control.

## Supporting Information

**S1 Fig.**
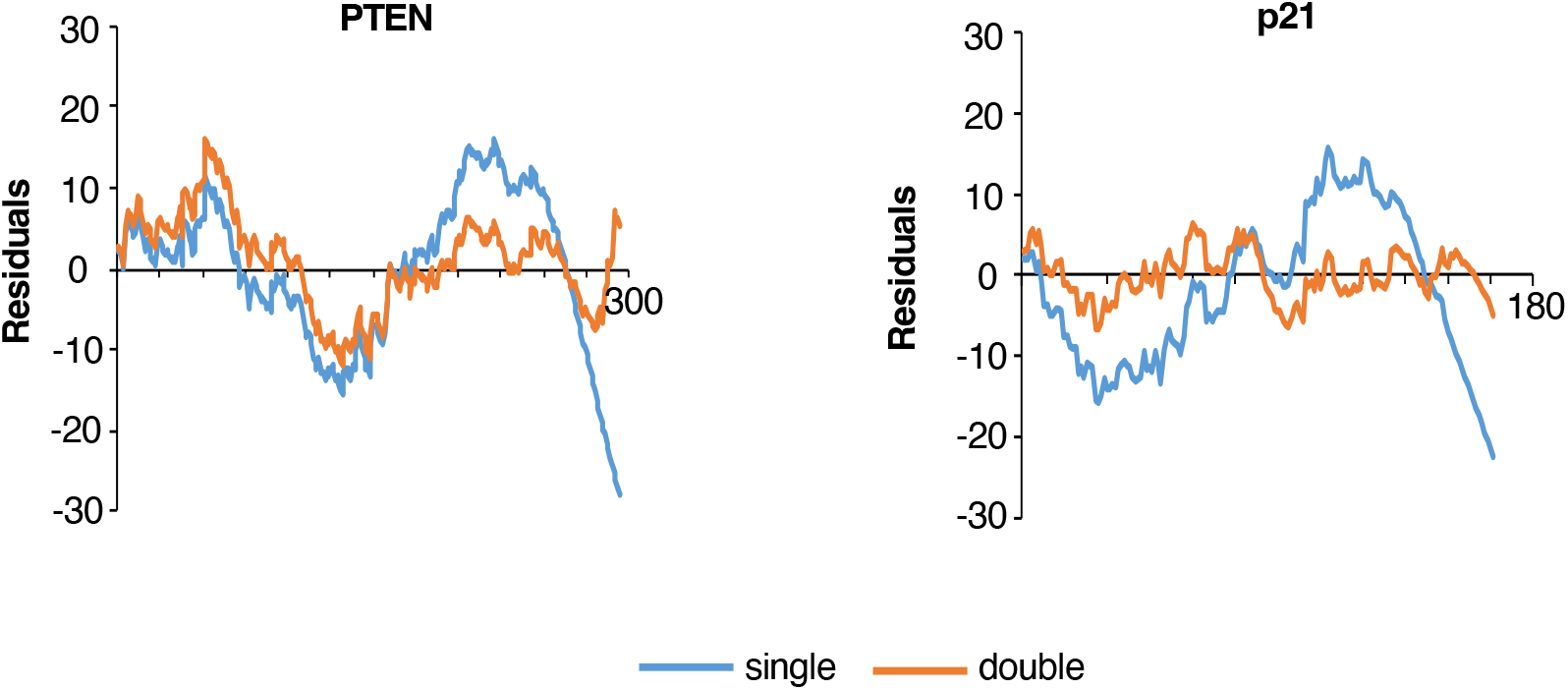
Plots of residuals from the exponential fits of the unbound dwell times for PTEN and p21 REs. The blue lines are from the single exponential fits and the orange lines are from the double exponential fits.

**S2 Fig.**
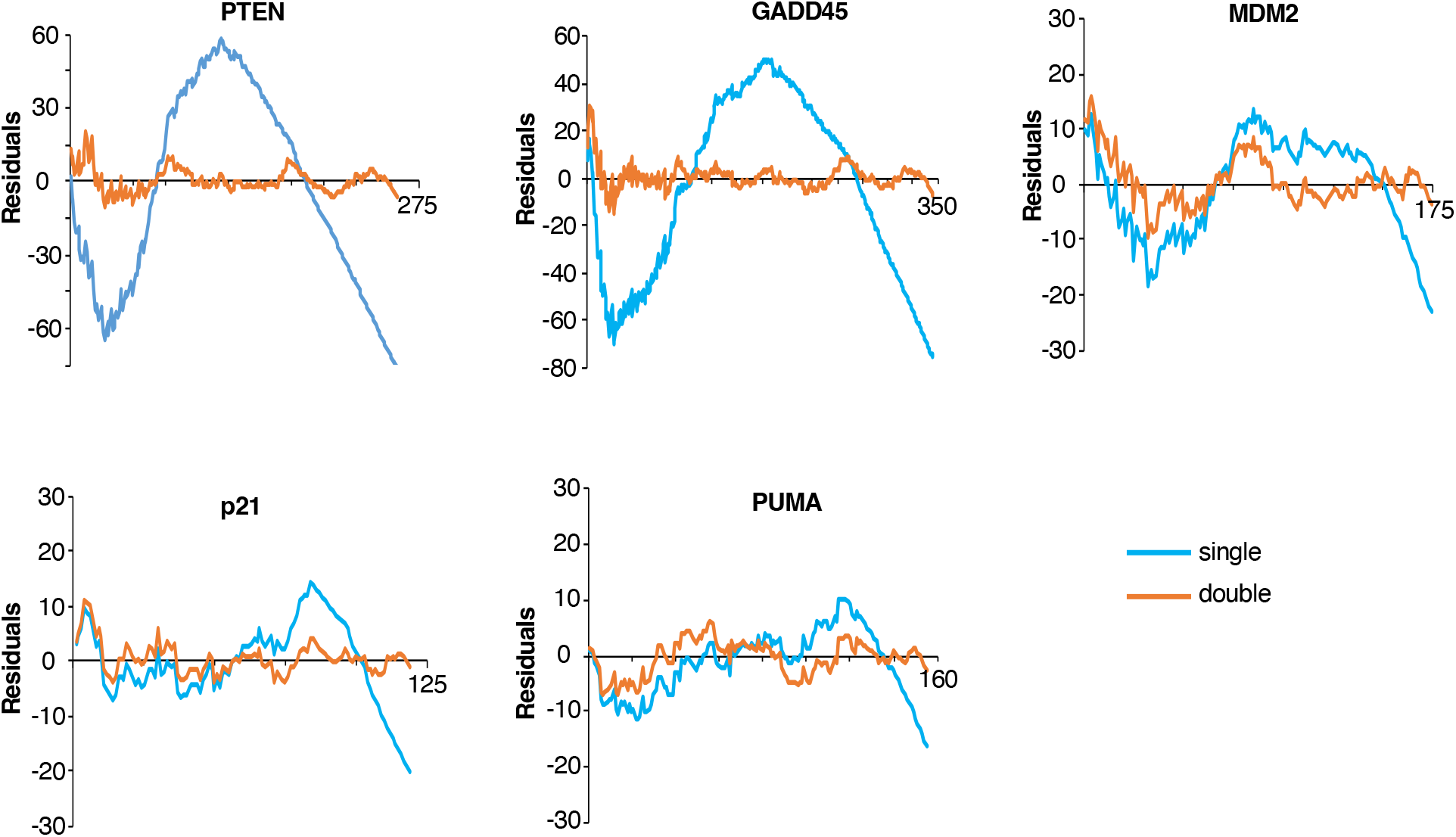
Plots of residuals from the exponential fits of the unbound dwell times. The blue lines are from the single exponential fits and the orange lines are from the double exponential fits. For all five REs the double exponential is a better fit for the data.

